# Inferring directional relationships in microbial communities using signed Bayesian networks

**DOI:** 10.1101/2020.02.18.955344

**Authors:** Musfiqur Sazal, Kalai Mathee, Daniel Ruiz-Perez, Trevor Cickovski, Giri Narasimhan

## Abstract

**Background:** Microbe-microbe and host-microbe interactions in a microbiome play a vital role in both health and disease. However, the structure of the microbial community and the colonization patterns are highly complex to infer even under controlled wet laboratory conditions. In this study, we investigate what information, if any, can be provided by a *Bayesian Network* (BN) about a microbial community. Unlike the previously proposed *Co-occurrence Networks* (CoNs), BNs are based on conditional dependencies and can help in revealing complex associations.

**Results:** In this paper, we propose a way of combining a BN and a CoN to construct a *signed Bayesian Network* (sBN). We report a surprising association between directed edges in signed BNs and known **colonization orders**.

**Conclusions:** BNs are powerful tools for community analysis and extracting influences and colonization patterns, even though the analysis only uses an abundance matrix with no temporal information. We conclude that directed edges in sBNs when combined with negative correlations are consistent with and strongly suggestive of colonization order.

## Introduction and Background

Bayesian Networks (BN) (also Belief Networks and Bayes Nets) are graphical models where nodes represent a set of multi-dimensional variables and edges represent *conditional dependencies* between the nodes. BNs can thus capture implicit and explicit relationships between these nodes [1]. When learning from data, edges in BNs can be directed or undirected. In fact, highly correlated variables very often lead to undirected (or two-way dependencies), since knowing one variable provides a lot of information about the other variable. In its simplest form, an edge in a BN expresses the conditional probability of knowing the (multi-dimensional) value of the variable at one node, given the value of the variable at another. BNs were used by Friedman et al. to use gene expression data to infer interactions between genes [2]. Conditional dependencies are often misinterpreted as *causation*, but are merely mathematical relationships that approximate causation under specific circumstances.

A significant feature of BNs is that they can allow us to differentiate between direct and indirect conditional dependence [3]. For example, if the dependence of variable *B* on variable *A* vanishes when conditioned on a third variable *C*, then it allows us to infer that a directed edge from *A* to *B* is superfluous and may be removed without loss of information since the directed edges (*A, C*) and (*C, B*) allows us to completely capture the dependency of *B* on *A*. BNs also help to differentiate between dependency configurations referred to as “common cause” and “common effect” [4].

Many algorithmic variants and implementations to construct BNs exist, including bnlearn [5], CG-BayesNet [6], Banjo [7], DEAL [8], GlobalMIT [9], BNFinder [10] and Tetrad [11].

Causation is an important type of relationship to be explored with biological data. So it makes sense to see if BNs can identify relationships that are suggestive of causation and that could lead to wet lab experiments for validation. Recently, BNs were used by Zhang et al. to understand changes in gene regulatory networks under different cellular states [12]. By modeling metabolic reactions and their involvement in multiple subnetworks of “metabosystems”, Shafiei et al. used BNs to infer differential prevalence of metabolic subnetworks within microbial communities [13].

The term *microbiota* refers to the community of microbes, including bacteria, archaea, protists, fungi, and viruses that share an environmental niche [14]. The term *microbiome* refers to the entire habitat, including the microbes, their genetic material and the environmental factors. The total genome from microbiota is referred to as the *metagenome*. The microbes exist as a *social network* because of the complex set of potential interactions between its various taxonomic members [15,16].

To understand potential interactions between taxa in a microbial community, the construction of co-occurrence networks (CoN) was proposed by Fernandez et al. [15] and Faust et al. [17]. The results suggested that groups of taxa frequently co-infected or co-avoided cohorts of subjects due to underlying interactions between them. Unfortunately, that is as far as CoNs are able to go in terms of inferring complex relationships in microbiomes.

In this paper, we investigate how to infer directional relationships between microbial taxa in a microbiome by focusing on the important challenge of inferring “colonization order” from abundance data.

In humans, normal microbial colonization starts from birth, and over time these communities become relatively stable [18]. Microbial communities are dynamic, and their compositions change with time [19]. Some microbes occupy an environmental niche early and then recruit other microbes suggesting an order of colonization in many microbial communities. Once new recruits enter the scene, their fitness for the environmental niche could determine the growth or decline of the early colonizers [20].

In the healthy state, our bodies harbor rich communities of microbes mostly on cutaneous and mucosal surfaces such as the skin, oral cavity, gastrointestinal tract, and the reproductive tract [21, 22]. Microbes in these communities have a variety of interactions that impact the health of the host or the environmental niche [17]. An imbalance (dysbiosis) in the microbial community is strongly associated with a variety of human diseases [23]. The dysbiosis is often due to invasion or increase in harmful pathogenic bacteria, which in turn is often preceded by colonization at the site of infection by specific early colonizers [24]. Thus, understanding colonization and its order can provide a window into how infections take hold. Understanding these functional (directed) relationships within the niche is critical for understanding healthy versus diseased microbiomes as well as the mechanisms and biological processes involved in the disease.

In this paper, we show that *signed Bayesian Networks* (sBNs), a variant of BNs obtained by combining BNs with *co-occurrence networks* can help tease apart some of these directed relationships and provide a glimpse into the complex and dynamic world of microbial communities. The paper is organized as follows. Section 2 provides foundations of BNs and some background on microbial colonization in select niches. Section 3 presents the details of the data and experiments and summarizes the results, and Section 4 presents some conclusions and future directions.

## Methods

### Bayesian networks

*Bayesian Networks* (BNs) are a class of *Probabilistic Graphical Models* (PGMs) [1,25] where each node represents a random variable from a set, **X** = {*X*_*i*_, *i* = 1, …, *n*}, with *n* random variables. The BN is represented as a graph *G* = (*V, E*), where each vertex in *V* represents a random variable from **X**, and *E* is the set of edges on *V*. In general, a BN is represented as a Directed Acyclic Graph (DAG), although undirected edges are used in cases where the direction cannot be reliably determined or when both directions appear plausible. Each random variable *X*_*i*_ has a local probability distribution. A directed edge of *E* between two vertices represents direct stochastic dependencies. Therefore, if there is no edge connecting two vertices, the corresponding variables are either marginally independent or conditionally independent given (a subset of) the rest of the variables. The “local” probability distribution of a variable *X*_*i*_ depends only on itself and its parents (i.e., the vertices with directed edges into the node *X*_*i*_); the “global” probability distribution, *P* (**X**) is the product of all local probabilities, i.e., a joint distribution [26] as shown below:

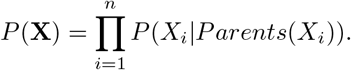

The task of fitting a BN is called “model learning” and its implementation generally involves two steps - *structure learning* and *parameter learning*. Structure learning involves finding a BN that encodes the conditional dependencies from the data, while parameter learning is the estimation of the parameters of the global distribution [27]. Eliminating edges in the structure helps to simplify the “global” joint distribution, allowing for more efficient computations with the model and for better inferring of critical relationships. In this paper, we only focus on the structure of the BN, not the parameters. For structure learning, at least three approaches have been proposed in the literature – constraint-based, score-based, and hybrid. We focus on the constraint-based algorithms, which are based on an approach called *Inductive Causation* (IC) [28]. IC provides a framework for learning the BN using conditional independence (CI) tests under the assumption that graphical separation in the BN is equivalent to probabilistic independence between the corresponding variables. Note that the resulting BN may be a partially directed acyclic graph (PDAG) [29] because not all edge directions can be resolved with IC.

### Training the Bayesian network structure

The constraint-based IC approach to structure learning mentioned above was proposed by Spirtes et al. [30]. The constraint-based approaches are typically more conservative than score-based algorithms in terms of the number of edges they retain in the final Bayesian network. Furthermore, constraint-based approaches are better suited for causal inferences [29]. The approach of Spirtes et al. was later modified by Colombo and Maathuis to make it *order independent* in an algorithm known as *PC-Stable* [31]. The main feature of PC-Stable algorithm is the inference of a skeleton (undirected structure) in an order independent way [31]. Order dependency is a minor issue for low dimensional settings. However, in high dimensional settings, order dependence may give results with high variance [32].

PC-stable consists mainly of three steps – adjacency search in order to learn the “skeleton”, identifying important substructures called *v***-structures**, and detecting and orienting other arcs. In Step 1, the algorithm starts with a complete undirected graph and then performs a series of conditional independence tests to eliminate as many edges as possible. The remaining undirected graph is referred to as the *skeleton*.

Step 2 is key to inferring a BN model, and uses the concept of *v*-structures, which are defined as follows. For any three nodes representing variables *X*_*i*_, *X*_*j*_, *X*_*k*_ in a Bayesian network *G*, if {*X*_*i*_, *X*_*j*_} and {*X*_*j*_, *X*_*k*_} are edges in *G*, but {*X*_*i*_, *X*_*k*_} is not, and if edges are oriented as *X*_*i*_ → *X*_*j*_ ← *X*_*k*_ then the triple (*X*_*i*_, *X*_*j*_, *X*_*k*_) is called a *v*-structure. Triples satisfying the *v*-structure property can be identified in the skeletons using conditional dependency tests, following which edges are appropriately directed to form a *v*-structure. The variable *X*_*j*_ in the triple forming the *v*-structure represents a “common effect” of *X*_*i*_ and *X*_*k*_. These *v*-structures are critical in giving directions to some of the edges of the skeleton.

In Step 3, three rules [31] are applied repeatedly to orient edges not already in *v*-structures.

**Rule 1:** Orient *X*_*j*_ − *X*_*k*_ as *X*_*j*_ → *X*_*k*_ whenever (a) there is a directed edge *X*_*i*_ → *X*_*j*_ and (b) *X*_*i*_ and *X*_*k*_ are not adjacent.

**Rule 2:** Orient *X*_*j*_ − *X*_*k*_ as *X*_*j*_ → *X*_*k*_ whenever there is a chain *X*_*j*_ → *X*_*i*_ → *X*_*k*_.

**Rule 3:** Orient *X*_*j*_ − *X*_*k*_ as *X*_*j*_ → *X*_*k*_ whenever there are two chains *X*_*j*_ −*X*_*i*_ → *X*_*k*_ and *X*_*j*_ −*X*_*l*_ → *X*_*k*_ given that *X*_*i*_ and *X*_*l*_ are not adjacent.

### Real Data sets

Ribosomal 16S rRNA sequences from three microbiome data sets (oral, infant gut, and vaginal) were used (see Table 1). The oral data set was generated as part of the Human Microbiome Project (HMP) from eight different sites within the oral cavity from 242 healthy adults (129 males, 113 females) [14, 33]. The samples included: saliva, buccal mucosa (cheek), keratinized gingiva (gums), palatine tonsils, throat, tongue dorsum, and supra- and sub-gingiva dental plaque (tooth biofilm above and below the gum) [14, 33].

**Table 1:**
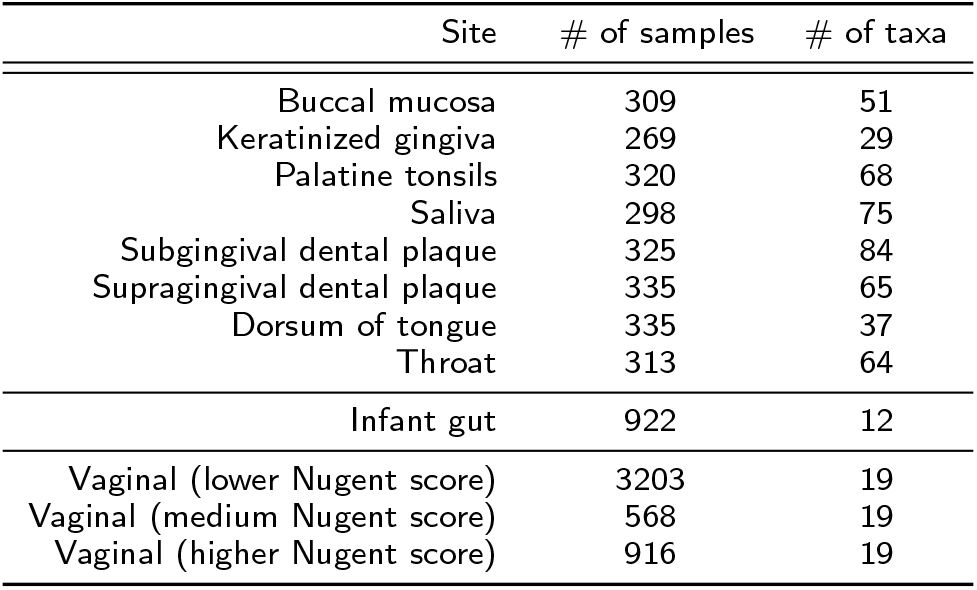
Microbiomes analyzed with sites, number of samples and number of taxa detected. The first eight are from oral microbiomes, the next one from gut microbiome, and the last three from vaginal microbiomes. Note that the Nugent score is an indicator of the level of vaginosis.

The preterm infant gut microbiome samples were collected and processed for a longitudinal study as described by La Rosa et al. [34]. This study involved a total of 922 stool samples from 58 premature babies, each weighing *≤* 1500 g at birth.

The vaginal microbiome data set was previously generated to determine temporal dynamics of the human vaginal microbiota [35]. This study involved 32 women from different ages (18 through 40), races (Black, White, Hispanic and other), educational backgrounds, and sexual habits [35]. Each sample was associated with a Nugent score [36], an indicator of the level of vaginosis. All OTUs associated with *Lactobacillus* were combined into one taxa.

Friedman et al. performed the BN inference by adding an extra “cell cycle phase” variable to account for the temporal aspect of the data [2]. Following their suggestion, an extra variable for sampling time was added to the analyses of the infant gut and vaginal microbiome data sets, thus assuming that the sampling time for each sample is an independent random variable from some distribution.

### Data processing

The samples were processed by amplifying the V35 hypervariable region of the bacterial 16S rRNA gene. This was followed by sequencing and grouping reads into common Operational Taxonomic Units (OTUs). The Mothur pipeline [37] was used to compute the microbial abundance of each taxon.

OTU abundance data were stored in matrix *B*, an *n × p* abundance matrix, where *n* is the number of samples and *p* is the number of OTUs. The *i*-th sample is represented by the *i*-th row of *B*, 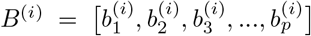, where 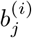 denotes the abundance of the *j*-th bacterial OTU in the *i*-th sample. The total number of mapped reads from the *i*-th sample is denoted by 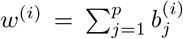. The relative abundance matrix is then computed by normalizing each raw count, 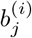, with the total number of reads in that sample *w*^*(i)*^. The normalized vector of relative abundances for sample *i* is thus given by

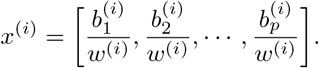

Each data set from the HMP collection had abundances for several hundred taxa, most of which were extremely small [14,33]. To make our computations efficient, taxa with abundance close to the background noise were eliminated. This is achieved by first sorting the relative abundance values of the OTU-level taxa and then picking the taxa with the highest values that added up to a total of 99%. In other words, the discarded taxa were the lowest values that summed up to less than 1%. Table 1 shows the number of taxa from each site used to learn the BNs during the structure learning step. The subjects in the vaginal data set were grouped by Nugent Scores — lower (healthy), medium, and higher. Individuals with higher Nugent scores had more severe cases of bacterial vaginosis [36].

### Semi-synthetic data

Besides using real data for our experiments, we also carried out experiments on what we refer to as “semisynthetic” data, which were obtained by appropriate modifications of real data sets as described below. The semi-synthetic data sets were obtained by performing temporal alignments on the infant gut data sets using the time-warping methods proposed by Lugo-Martinez et al. [38, 39]. The purpose of temporal alignments was to align the “internal clocks” of the subjects correcting for their different metabolic speeds. The temporal alignment was done by interpolating the time series and stretching/squishing and shifting them with respect to time series of a reference subject. As a consequence, the time series are put on an artificial time scale and then uniformly sampled with a sampling rate of 1 per (warped) day.

### Construction of Bayesian Networks

The PC-stable, a causality-learning algorithm, was used to construct the BNs [31]. It is a constraint-based algorithm that is more conservative than score-based algorithms and results in fewer false positives. Also, it is partly order-independent, as described below [31]. The PC-stable algorithm from the `bnlearn` package [40] was used to obtain the BNs for each data set.

### Construction of Co-occurrence Networks

The co-occurrence networks (CoNs) were constructed for each cohort using Pearson correlation coefficient, as described in previous work [15].

### Construction of Signed Bayesian Networks

The edges of BNs were augmented with the coefficient values generated in CoNs, thus distinguishing between positive and negative correlations. As mentioned earlier, the resulting network is referred to as a *Signed Bayesian Network* (sBN). All sBNs in this paper were visualized using Cytoscape [41]. The color of the edges (green for positive and red for negative) indicates sign information.

### Experiments and Statistical Analyses

The constraint-based algorithms employ statistical tests for deciding conditional independence. Since the random variables in our experiments hold continuous data representing the abundance of taxa, we used linear correlation (*student’s exact T-test*) and Fisher’s Z-test (*asymptotic normal test*) for conditional independence testing [42, 43].

In the PC-stable algorithm, inferring the skeleton structure and inferring the directions of edges involved in the *v*-structures are known to be “order-independent”. However, inferring the directions of edges not involved in the *v*-structures is not order-independent. A non-parametric bootstrap value was computed to indicate the strength of each edge in the output network in order to assess the accuracy of the output [44, 45]. To achieve this, the data was randomized before input into the PC-stable algorithm. The bootstrap values were computed by executing the program on 200 different permuted inputs and reporting the percentage of times it reports one direction.

## Results and Discussion

The sBNs were obtained by prudent use of BNs in conjunction with CoNs. The main contribution of this paper is to show evidence to support the claim that sBNs can help make inferences about *colonization order*. In some niche environments, research has shown that microbes colonize the niche in specific orders, with early colonizers often recruiting late colonizers or creating conditions that make it more attractive for specific late colonizers [46]. We have observed that with high accuracy, the edges of sBNs are consistent with known colonization orders. In particular, we show that the sBNs can capture colonization order when augmented with the correlation coefficient. The findings were validated by analyzing oral, infant gut, and vaginal microbiome data sets, where prior published information on colonization order was available. The colonization order was also retained in our experiments with the semi-synthetic data sets as well.

The sBNs generated from the data sets mentioned above were visualized with Cytoscape. In all the sBNs generated (Figs. 2–5 and Supplementary Figs. 6-11), nodes correspond to bacterial taxa, node sizes are proportional to the average abundance of the taxa, thickness of the edges are proportional to the absolute value of Pearson correlation coefficient (i.e., measure of co-occurrence), and opacity of an edge is proportional to its bootstrap values. Edges are colored green and red for positive and negative correlations, respectively. The purple and red node colors correspond to the bacterial taxa that are described as early and late colonizers (in published literature), respectively [47–49]. The black nodes indicate colonizers whose order has not been described previously. We note (data not shown) that while there are many strongly connected clusters in CoNs, these nodes remain connected in sBNs (as expected), but relatively sparsely because of the stringent conditional probability tests.

### Semi-synthetic data from infant gut microbiome — sBN Edges are Consistent with Temporal Order

The infant gut data set was temporally aligned as described earlier. We then divided the time line into *k* periods, with *k* = 1, 2,... and created sBNs from each period. The goal was to see if any of the known orders of colonization can be observed in the figures, even after having modified the time axis of each subject differently.

The infant gut is dominated by three classes that generally appear and colonize in a sequential order: Bacilli (Firmicutes) soon after birth, which then gives way to the Gammaproteobacteria (Proteobacteria), and followed by Clostridia (Firmicutes) [34]. When we partitioned the time series into *k* = 2 periods, the sBN from the first period had a directed edge from the Bacilli to Gammaproteobacteria. The red-colored edge suggested a negative correlation as would be expected if this inference came from colonization order. Additionally, the sBN generated from the second period showed a directed edge from Gammaproteobacteria to Clostridia, also colored red (Fig. 1).

**Figure 1:**
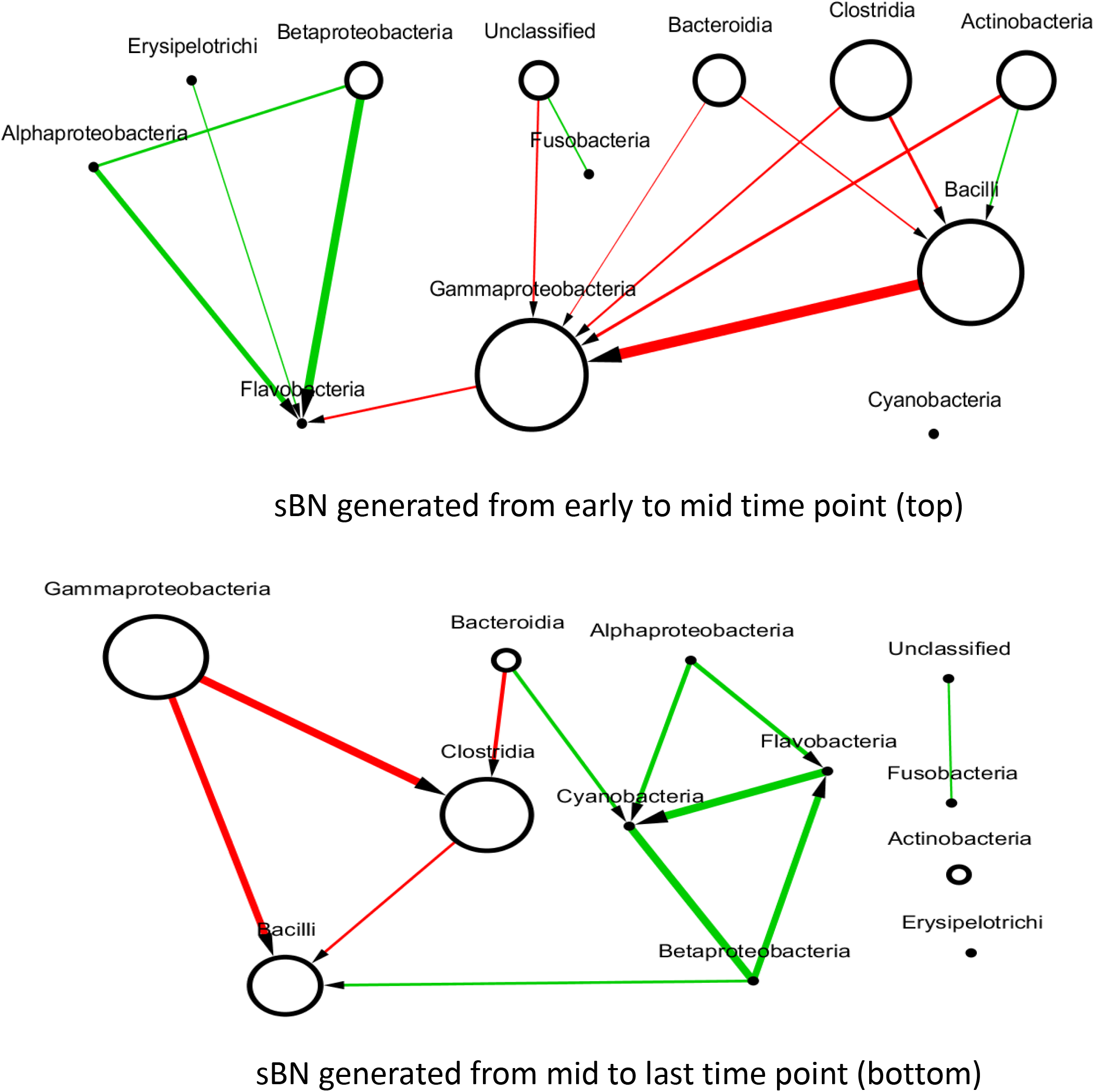
sBN of semi synthetic infant gut microbiomes

When the time series were partitioned into three periods, the same two edges were represented strongly in periods 2 and 3 respectively. In fact, the strength of the two edges in the three periods were (1) 0.4 and 0.16 (i.e., both weak), (2) 0.94 and 0.16, and (3) 0.61 and 0.80. The above observations suggest strongly that the transition from Bacilli to Gammaproteobacteria occurs before the transition from Gammaproteobacteria to Clostridia, and that the colonization order is supported in the sBNs.

We, therefore, conclude that sBNs are capable of capturing colonization order using the methods suggested above. Red edges or negative correlations are consistent with the model that for both edges when one taxon is declining in abundance, the other is increasing in abundance.

### Oral Microbiome — sBN Edges are Consistent with Colonization Order

In the oral cavity, early and late bacterial colonizers have been identified and reviewed in the literature [47]. Many species from the genus *Streptococcus* is the early primary colonizer, accounting for 60% 90% of the early abundance profile [50]. The following taxa have been identified as early and late colonizers for oral microbiomes [47–49].

**Early:** *Streptococcus gordonii*, *Streptococcus mitis*, *Streptococcus oralis*, *Streptococcus sanguis*, *Actinomyces israelii*, *Actinomyces naeslundii*, *Propionibacterium acne*.

**Late:** *Selenomonas flueggei*, *Treponema* spp., *Porphyromonas gingivalis*.

Comparison of the sBNs for all oral microbiomes (Figs. 2–3 and Supplementary Figs. 6-11) showed that the keratinized gingiva (Fig. 2) and tongue dorsum (Fig. 3) have the fewest number of distinct taxa. The sBNs for these two sites were more distinctive than those derived from other sites and showed stronger correlations between taxa. The saliva, subgingival, and palatine tonsils sites harbored a higher number of taxa and exhibited weaker correlations. Note that not every taxa is present in every oral site, thus explaining the differences in the set of nodes present in each sBN.

**Figure 2:**
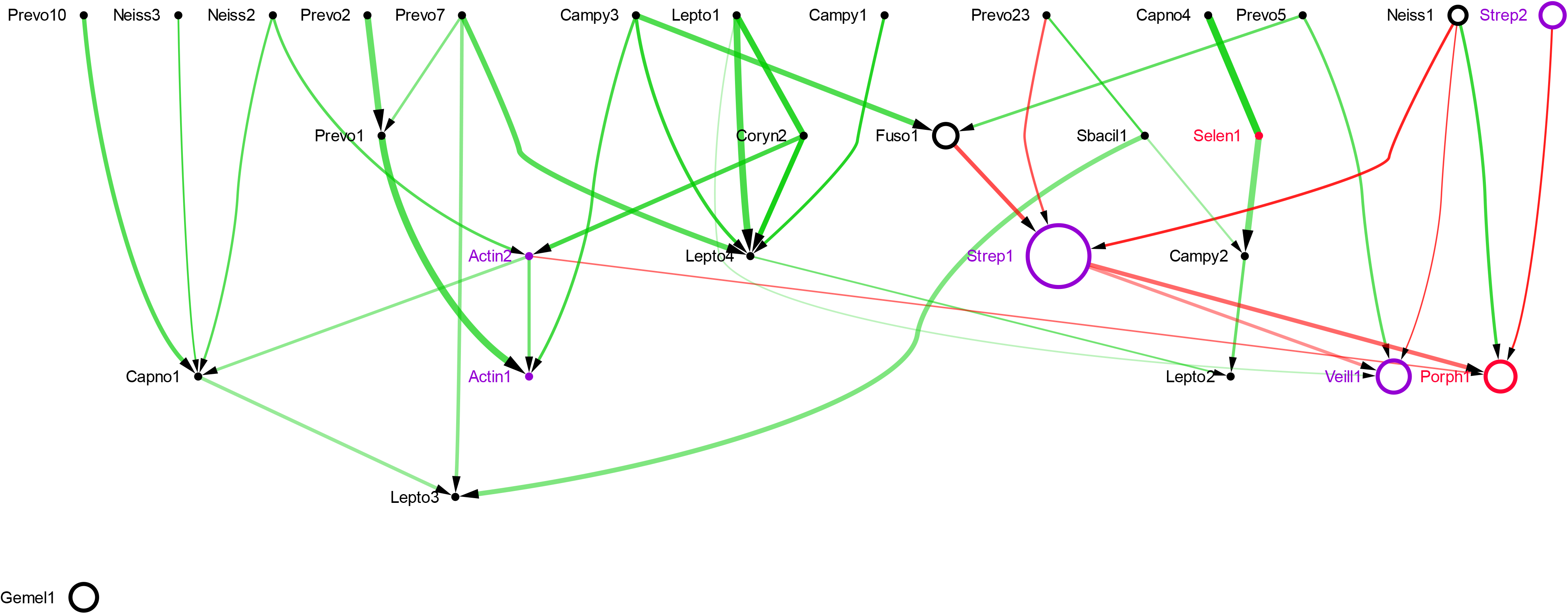
sBN of Keratinized gingiva

**Figure 3:**
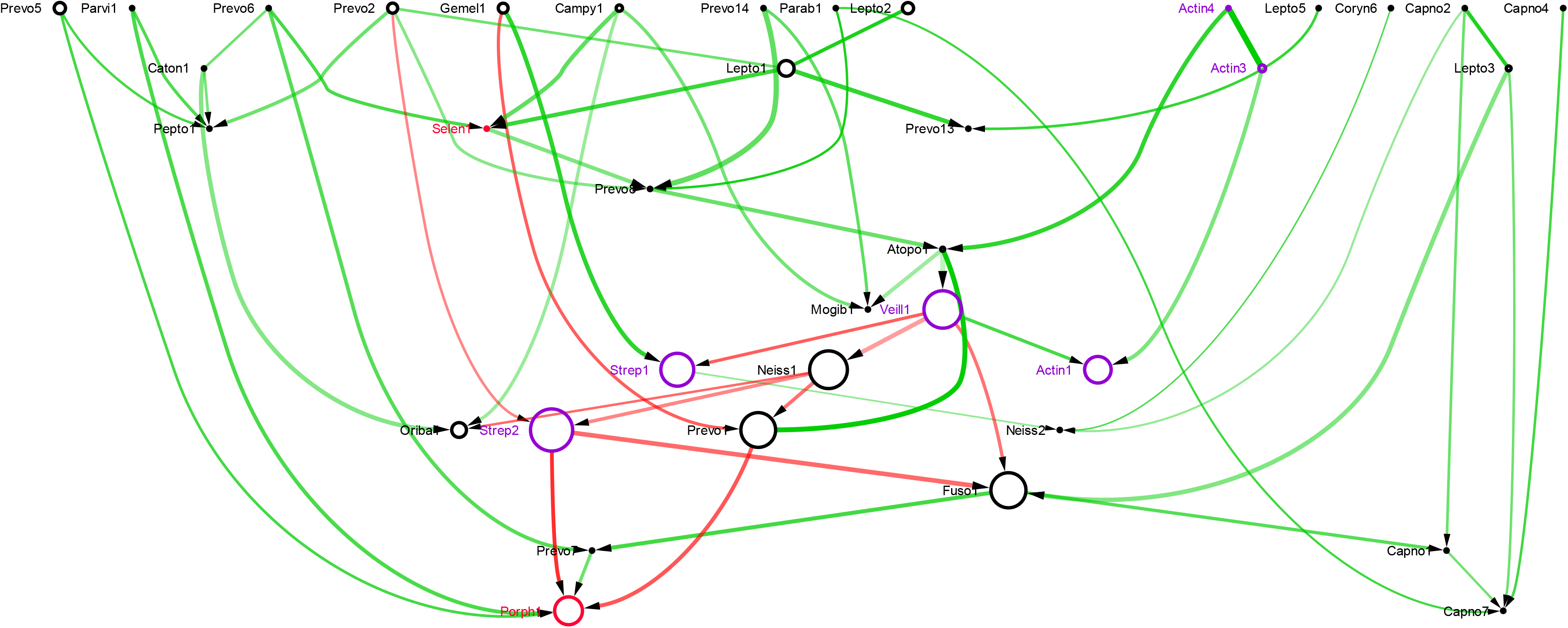
sBN of Tongue dorsum

The sBNs for the oral microbiomes had a combined total of 716 edges. Of these, 78 edges connected vertices, which were associated with known early or late colonizers. Table 2 summarizes the directed edges between early and late colonizers, they are consistent with the known colonization order, and the correlation (negative/positive edges) among them. More than 90% of the sBN edges for the oral microbiome were directed with the exceptions of saliva and buccal mucosa, for which only 83-84% were directed. Of the 78 edges connecting labeled vertices, all edges except for two were consistent with the known colonization order, i.e., directed from early to late colonizers (Table 2). These two edges are shown as dashed lines in the corresponding sBNs (see Figs. 7 and 10). In summary, for the oral microbiome the directed sBN edges go from early to late colonizers, with few exceptions. For example, the sBN from keratinized gingiva (Fig. 2) has three directed edges (Actinomyces2-Porphyromonas1, Streptococcus1-Porphyromonas1, and Streptococcus2Porphyromonas1) from early colonizers to late colonizers and none from late to early colonizers. Note that all taxonomic names have been abbreviated in the figures to the first five characters plus a number, each name refers to a distinct OTU. The sBN for the buccal mucosa (Supplementary Fig. 6), palatine tonsils (Supplementary Fig. 7), saliva (Supplementary Fig. 8), subgingival plaque (Supplementary Fig. 9), supragingival plaque (Supplementary Fig. 10), and throat (Supplementary Fig. 11) are included in the supplementary files.

**Table 2:** Inferring Colonization order in oral microbiomes. The columns indicate the following: sampled oral sites, total number of edges in causal network, number of directed edges, total number of negatively correlated (red) edges, number of edges connecting early to late colonizers, number of edges connecting early with early or late colonizers, number of inconsistent directed edges (i.e., from late to early colonizers), and percentage of negatively correlated edges connecting early to late colonizers.

| Oral Site | Total | Directed | Red | $E \rightarrow L$ | $E \rightarrow E$ or $E \rightarrow L$ | $L \rightarrow E$ | Consistent Red edges |
| --- | --- | --- | --- | --- | --- | --- | --- |
| Buccal mucosa | 69 | 57 | 5 | 1 | 4 | 0 | 100% |
| keratinized gingiva | 39 | 36 | 8 | 3 | 5 | 0 | 100% |
| Palatine tonsils | 126 | 116 | 9 | 1 | 13 | 1 | 92% |
| Saliva | 102 | 86 | 8 | 1 | 12 | 0 | 100% |
| Subgingival plaque | 123 | 113 | 8 | 1 | 18 | 0 | 100% |
| Supragingival plaque | 109 | 105 | 11 | 3 | 13 | 1 | 92% |
| Dorsum of tongue | 56 | 50 | 11 | 1 | 4 | 0 | 100% |
| Throat | 92 | 83 | 8 | 0 | 9 | 0 | 100% |
| Total | 716 | 646 | 68 | 11 | 78 | 2 | 97.4% |

### Oral Microbiome — sBN Edges with Negative Correlation are Consistent with Colonization Order

As mentioned above, two out of the 78 edges are exceptions to the rule that no edges in the sBNs are directed from late to early colonizers. In particular, one edge goes from Trepo5 (*Treponema*, labeled as a late colonizer) to Actin3 (*Actinomyces*, labeled early colonizer) in palatine tonsils. Similarly, another edge goes from Porph3 (*Porphyromonas*, labeled as late colonizer) to Actin3 (*Actinomyces*, labeled early colonizer) in supragingival plaque. However, the correlation coefficient of the edges between them is positive. Thus, the accuracy in terms of direction is 97.4%, and all correctly directed edges have negative correlations. According to Kolenbrander et al., the bacterial taxa representing early colonizers coaggregate with only a specific set of other early colonizers, and not with any of the late colonizers [47]. Our findings, albeit limited, are consistent with this observation, that all edges connecting early to late colonizers in that direction are negatively correlated (red edges).

### Infant Gut Microbiome

The abundance of microbes in neonatals over the course of the first few weeks of their lives have been reported [34]. In two infant gut microbiome studies, the class Bacteroidetes and Gammaproteobacteria were observed early, followed by Bacilli, Clostridia and Gammaproteobacteria [34, 51]. Over time, there was a significant decrease in Bacilli, and the infant’s gut appears to have a tug-of-war between the two classes Gammaproteobacteria and Clostridia [51]. When the sBNs were constructed with the infant gut microbiome data, we obtained a directed network that supported the claim that sBNs shed light on the colonization pattern (Fig. 4). There were directed edges from Bacteroidetes, Bacilli, and Clostridia to Gammaproteobacteria (Fig. 4). The results also supported the prior knowledge that Clostridia precedes Bacilli in the colonization order. All these taxa are mostly negatively correlated (red edges), as shown in Fig. 4, reinforcing the point that a directed edge combined with negative correlations is strongly suggestive of colonization order.

**Figure 4:**
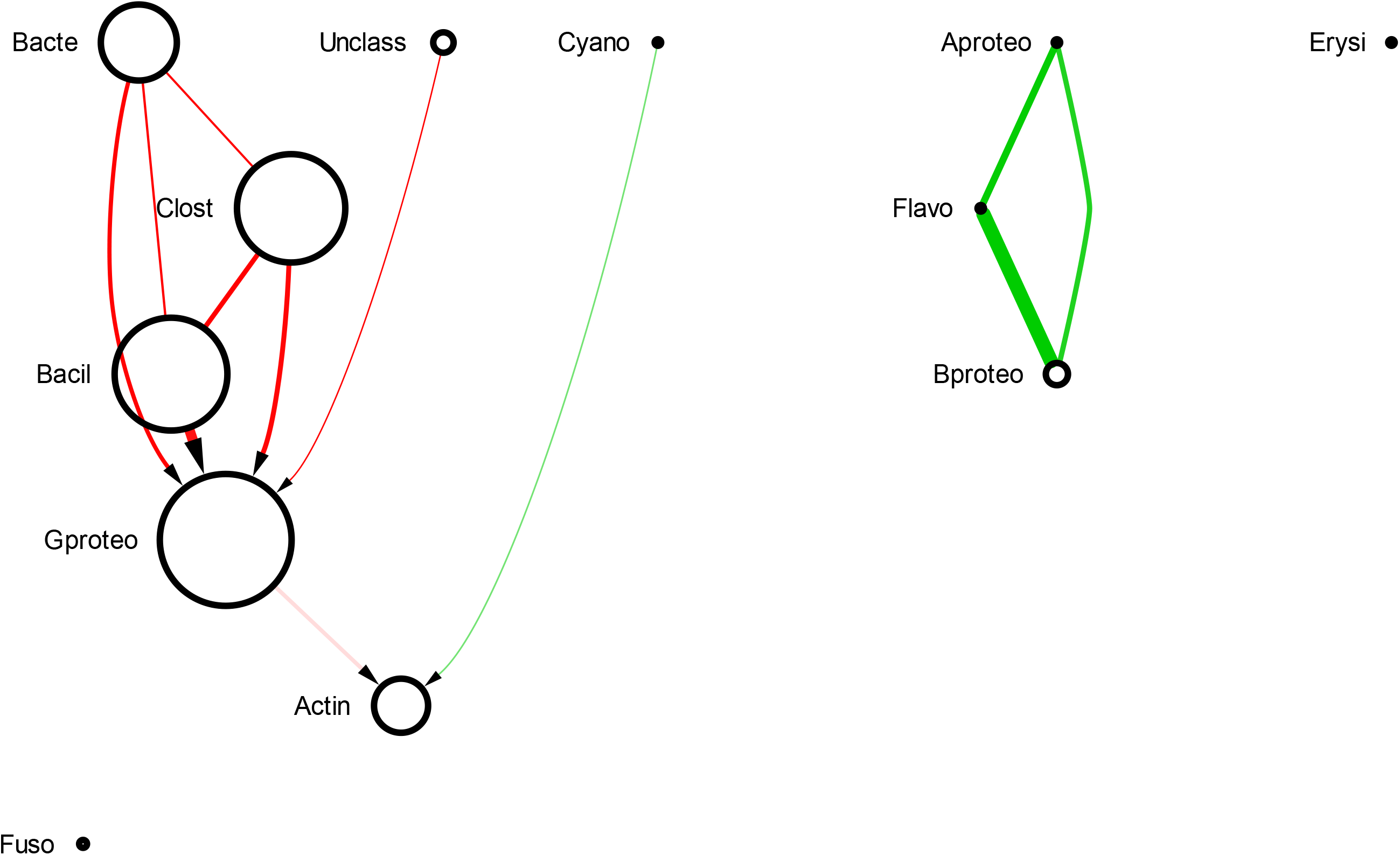
sBN of infant gut microbiome

### Vaginal Microbiome

A healthy vaginal microbiome is dominated mainly by *Lactobacillus* species [52]. When women at a reproductive age suffer from bacterial vaginosis (BV), the *Lactobacillus* species are replaced by *Gardnerella*, *Peptostreptococcus*, *Atopobium*, *Sneathia*, *Parvimonas*, and *Corynebacterium*, among others [53]. Fig. 5 shows three sBNs for vaginal microbiomes associated with low (healthy), medium (early BV), and high (advanced BV) Nugent scores. All samples were analyzed for the abundance of the same set of 23 genera. Overall, the predominant genera observed were *Lactobacillus*, *Atopobium*, *Gardnerella*, *Parvimonas*, and *Prevotella* (Fig. 5).

**Figure 5:**
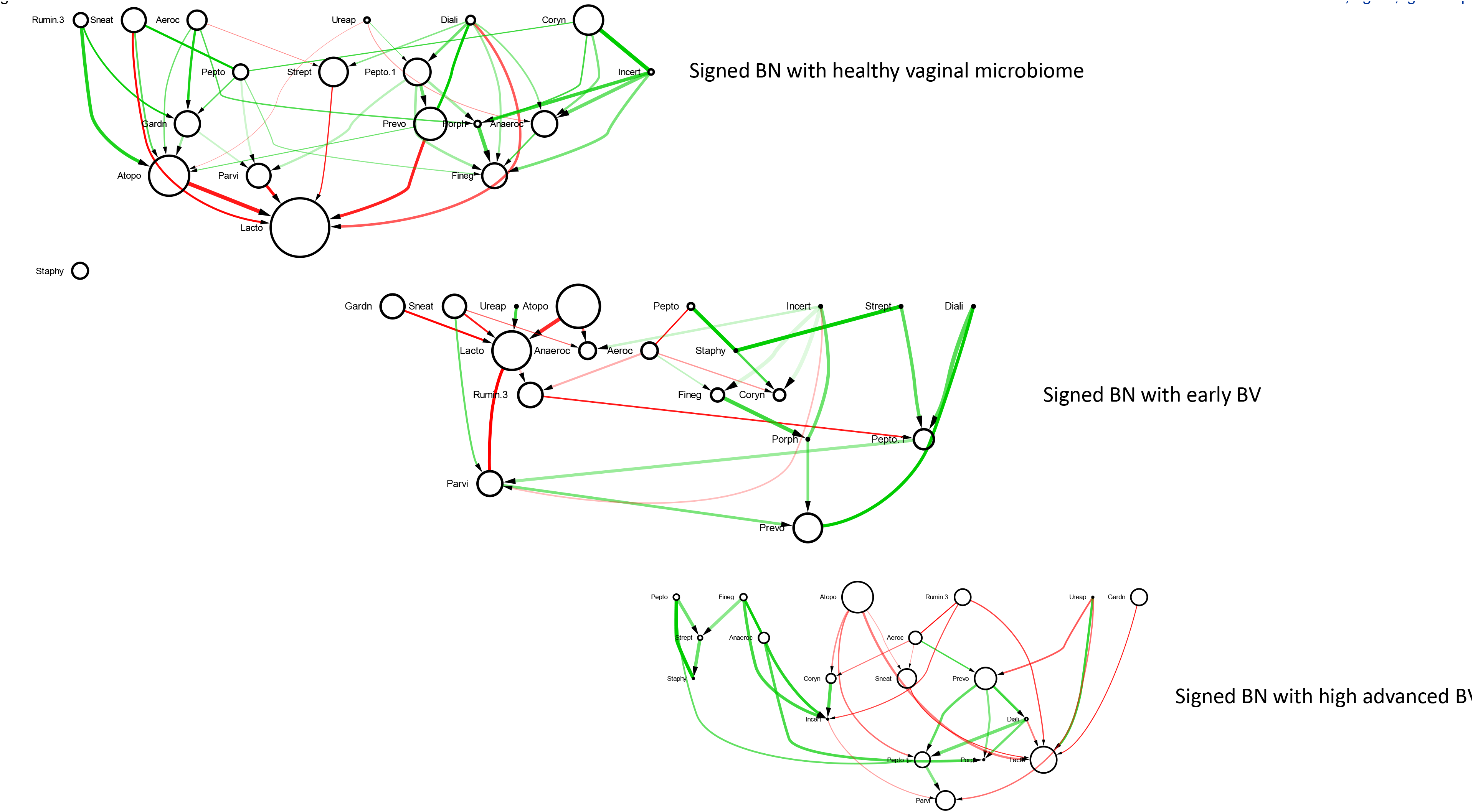
sBN of vaginal microbiome

In the sBN associated with the healthy “vaginome”, the abundance of *Lactobacillus* was comparatively higher as expected. The *Lactobacillus* species, especially, *L. crispatus* and *L. iners* (data not shown) displayed an antagonistic relationship with the BV-associated *Gardnerella*.

In the sBN for the medium Nugent score cohort, indicative of early vaginosis, the BV-associated genera, *Atopobium*, and *Sneathia* AND *Gardnerella* were significantly increased in abundance, and appeared as early colonizers. The abundance of all the BV-associated pathogens was negatively correlated with *Lactobacillus*, reaffirming an antagonistic relationship. In the sBN for the advanced BV cohort, characterized by higher Nugent scores, a proportional increase in abundance was observed with *Atopobium* followed by *Gardnerella*. Even with the antagonistic relationship with *Lactobacillus*, the BV-associated pathogenic genera especially *Atopobium* and *Gardnerella*, *Sneathia* are connected by a directed edge to *Lactobacillus*. The appearance of the pathogenic genera as late colonizers is consistent with clinical findings [54]. Strong positive relationships were observed between *Prevotella* and *Peptostreptococcus*, and *Peptostreptococcus* with *Parvimonas*. This may suggest that the presence of *Prevotella* enables the colonization of *Peptostreptococcus* followed by *Parvimonas*.

To check the robustness we also experimented with a higher number of taxa, i.e., by including all taxa whose abundance added up to 99.99 %. We found that sBNs can retrieve the known colonization order even if we include taxa with small abundance (from 99% to 99.99% of most abundant taxa shown in Fig. 12).

## Conclusions

In healthy oral microbiomes, taxa such as *Actinobacteria* were identified as early colonizers [55]. Many pathogenic microbes associated with oral diseases such as dental caries, gingivitis, and periodontitis appeared as late colonizers [56]. In addition, there were antagonistic relationships between these pathogens. The rivalry seemed to occur between *Streptococcus*, *Fusobacterium, Prevotella, Porphyromonas, Veillonella, Propionibacterium* and *Neisseria*. Since the oral samples came from healthy individuals, the existence of the rivalry could lead to the elimination of one or more taxa from the site. Alternatively, it is also possible that one taxon keeps the other in check to prevent dysbiosis. A well-known pathogenic genera, *Treponema*, appeared as a late colonizer with positive correlations in most of the sites. It was absent in keratinized gingiva and tongue dorsum, but appeared as an early colonizer in buccal mucosa. This may suggest that the buccal mucosa is the site in the oral cavity where *Treponema* colonizes.

The sBN for the vaginal microbiome confirmed previously known relationships between *Lactobacillus* and other BV-associated pathogens. In the process, it also suggested a possible colonization order. It would require a longitudinal study of women before and after BV to validate the suggested colonization order. Current analyses suggest that the balance in the relative abundance of *Lactobacillus* and *Atopobium* may be a *biomarker* for BV.

Inferring the interactions between different taxa within a microbial community and understanding their influence on health and disease is one of the primary goals of microbiome research. The sBNs help us to infer potential relationships and dependencies within a microbiome, and the colonization order, even without the use of data from longitudinal studies. The sBNs could help in understanding the dependencies between the entities of a microbial community.

Finally, we reiterate the conclusion that directed edges in sBNs when combined with negative correlations, may be strongly suggestive of colonization order.

## List Of Abbreviations

BN: Bayesian Network
CoN: Co-occurence Network
IC: Inductive Causation
CI: Conditional Independence
PDAG: Partially Directed Acyclic Graph
PGM: Probabilistic Graphical Model
OTU: Operational Taxonomic Unit
HMP: Human Microbiome Project
sBN: Signed Bayesian Network
BV: Bacterial vaginosis

## Declarations

### Ethics approval and consent to participate

Not applicable.

### Consent for publication

Not applicable.

### Availability of data and material

Raw data for the oral microbiome dataset were downloaded from the Human Microbiome Project website (https://hmpdacc.org/hmp/). The infant gut microbiome dataset was downloaded from the supplementary materials of La Rosa et al. [34]. The vaginal microbiome dataset was downloaded from the supplementary materials of Gajer et al. [35]. Source code and processed data can be available upon reasonable request to the corresponding author.

### Author’s contributions

The research project was conceived and supervised by GN. MS wrote all necessary software and performed all the experiments. DR and TC assisted with data processing. KM contributed to writing and interpreting biological significance. All authors reviewed the manuscript.

## Acknowledgments

The authors thank the members of the Bioinformatics Research Group (BioRG) for many useful comments during the course of this research.

---

Figure 6: (Suppl.) sBN of Buccal mucosa

Figure 7: (Suppl.) sBN of Palatine tonsils

Figure 8: (Suppl.) sBN of Saliva

Figure 9: (Suppl.) sBN of Subgingival plaque

Figure 10: (Suppl.) sBN of Supragingival plaque

Figure 11: (Suppl.) sBN of Throat

Figure 12: (Suppl.) sBN of Keratnized Gingiva with 99.99 % taxa

